# *In vitro* screening of anti-viral and virucidal effects against SARS-CoV-2 by *Hypericum perforatum* and *Echinacea*

**DOI:** 10.1101/2021.01.11.426295

**Authors:** Leena Hussein Bajrai, Sherif Ali El-Kafrawy, Rabie Saleh Alnahas, Esam Ibraheem Azhar

**Affiliations:** Special Infectious Agent Unit, King Fahd Medical Research Center, King Abdulaziz University, Jeddah, Saudi Arabia.; Biochemistry Department, Faculty of Sciences, King Abdulaziz University, Jeddah, Saudi Arabia.; Department of Medical Microbiology and Parasitology, Faculty of Medicine, King Abdulaziz University, Jeddah, Saudi Arabia.

**Author notes:** conceived and designed the experiments, performed the whole experimental work, analyzed the data, and wrote the paper (Leena Hussein Bajrai). interpreted the data and reviewed the paper (Sherif Ali El-Kafrawy). edited the references and organized the light microscope photographs (Rabie Saleh Alnahas). co-corresponding author and reviewed the paper.

## Abstract

Special Infectious Agent Unit in King Fahd Medical Research Center at King Abdulaziz University, Jeddah, Saudi Arabia, has pursed the anti-viral project field to optimize the group of medicinal plants for human-infectious diseases. We have begun virtually in this field since COVID-19 pandemic, besides our divergence in the infectious agents’. In this study and based on the previous review, *Hypericum perforatum* (St. John’s Wort) and *Echinacea* (gaia HERBS^®^) were tested *in vitro* using Vero E6 cells for their anti-viral effects against the newly identified Severe Acute Respiratory Syndrome Coronavirus-2 (SARS-CoV-2) through its infectious cycle from 0 to 48 hours post infection. The hypericin (0.9 mg) of *H. perforatum* and the different parts (roots, seeds, aerial) of two types of *Echinacea* species (*Echinacea purpurea* and *Echinacea angustifolia*) were examined their efficacy in certain concentration and under light-dependent anti-viral activities to measure the inhibition of the SARS-CoV-2 mRNA expression of RNA-dependent RNA polymerase (RdRP) gene and the viral load with quantitative real-time polymerase chain reaction (qRT-PCR), and to assess the neutralization of the SARS-CoV-2 spike receptor binding on cell culture assay. Interestingly, the mixture (*H.E.*) of 100 mg/mL of *H. perforatum* and *Echinacea* was tested too on SARS-CoV-2 and showed crucial anti-viral activity competing *H. perforatum* then *Echinacea* effects as anti-viral treatment. Therefore, the results of gaia HERBS^®^ products, *H. perforatum* and *Echinacea* species, applied in this study showed significant anti-viral and virucidal effects in the following order of potency: *H. perforatum, H*.*E*., and *Echinacea* on SARS-CoV-2 infectious cycle; and will definitely required a set up of clinical trial with specific therapeutic protocol based on the outcome of this study.

**Author Summary:** After an outbreak of Rift Valley Fever in the Southern region of Saudi Arabia, particularly in May 2003, Special Infectious Agents Unit (SIAU) was established and founded by Prof. Esam Ibraheem Azhar. This unit contains a full range of facilities including Biosafety Level 3, allows him and his research groups to ambulate and culture risk group 3 viruses in Saudi Arabia & Gulf States for the first time. Since that time, SIAU and our international collaboration have been extended to implement a standard protocols in the infectious agents diagnostics procedure through different mode of collaboration including exchange of expertise, joint research program and more recently a technology transfer agreements with number of international institute sharing same interests. Furthermore, we have been engaged in number of researches related to Hajj & Umrah plus number of national services with the Ministry of Health (MOH) through which, we utilize our Mobile biosafety level 3 Lab to enhance the diagnostics of MERS CoV in the Holly sites during Hajj since 2014.

In our SIAU and with a powerful team, we have excellent researches made valuable contributions through *in vivo* and *in vitro* animal and human studies, and several human viral pathogens which are a threat to global health security due to millions of pilgrims visiting Saudi Arabia every year from 182 countries: with particular areas of interests in: Alkhurma Viral Hemorrhagic Fever, Dengue Hemorrhagic Fever Viruses, Rift Valley Fever Virus, MERS-CoV and more recently the new global infectious diseases threat, Sever Acute Respiratory Syndrome Coronavirus-2 (SARS-CoV-2).

## Introduction

Coronaviruses (CoVs) are a large group of RNA viruses with single-stranded RNA genomes that cause 30% of upper and lower respiratory tract infections in humans. Four main human coronaviruses (HCoVs) are known including severe acute respiratory syndrome SARS-CoV [1], HCoV-NL63 [2], HCoV-HKU1 [3], and MERS-CoV in 2012 [4]. SARS-CoV-2 originated from Wuhan, Hubei Province of China and spread to the rest of the world causing the global pandemic of COVID-19 with symptoms ranging from cough, fever, and dyspnea to neurological and vascular symptoms [5]. With advancement of the disease, intensive production of pro-inflammatory cytokines findings like: TNF-α, IL-1β, IL-6, IFN-γ, CXCL10, MCP-1, resulting in vasculitis, hyper-coagulability, and multi-organ damage which lead ultimately to death [6]. Main protease (Mpro), known as 3-chymotrypsin-like cysteine protease (3CLpro), has been considered as an important functional target in the viral life cycle, and therefore as a candidate target for anti-viral drugs against SARS-CoV-2 [7, 8] due to its role in the release of functional polypeptides that is encoded by all HCoVs [9, 10]. Updated reports about the disease management from COVID-19 have targeted its structure, pathology, and mechanism in order to have the best solution against the infection by both redesigning and using classical anti-inflammatory agents and by targeting certain cytokine of molecular drug, as per WHO recommendation [11]. For example, numbers of feasible treatment against SARS-CoV-2 were proposed such as: neuraminidase inhibitors, Remdesivir, Peptide (EK1), abidol, RNA synthesis inhibitors (TDF and 3TC), anti-inflammatory drugs (hormones and other molecules), and Chinese traditional medicine [12]. Furthermore, there were several studies about the synergy between antimalarial drugs, like Chloroquine-Hydroxychloroquine, Remdesivir-Favipiravir for the treatment of COVID-19 [13–15].

*H. perforatum* (family Hypericaceae), or St. John’s Wort (SJW) has been very well known for a long time as an effective medicinal plant for a range of communicable and non-communicable diseases such as depression, bacterial and viral infections, skin wound, and inflammation [16, 17]. Herbal medicine studies demonstrate the pharmacological efficacy against microbial infections [16, 18, 19]. As an anti-viral agent, *H. perforatum* activities were assessed *in vitro* and *in vivo* on infectious bronchitis virus (IBV), Hepatitis C, HIV and Coronaviruses other than SARS-CoV-2 [20, 21]. Several Studies showed its activity against *Escherichia coli, Shigella dysenteriae, Salmonell typhi, Bacillus cereus*, and several oral bacteria [22]. Because of *H. perforatum’s* metabolites from each extracted part (roots, seeds and aerial), they are chemically defined to naphthodianthrones (*hypericin*), phloroglucinols (*hyperforin*), flavonoid glycosides (*hyperoside*), biflavones, and anthocyanidins [19]. Therefore, the hyperforin and hypericin of *H. perforatum* are shown to be effective as antibacterial compounds against various Gram-positive bacteria but ineffective against Gram-negative bacteria [23–26]. Furthermore, ethyl acetate extraction section of Hypericum showed a significant reduction on the titer of IBV [16].

Another medicinal plant, which was applied for many traditional and common remedies like curing cold and flu symptoms and boosting immune system, is *Echinacea*. Echinacea is known with nine species of several plants in the genus of *Echinacea*, however, only three of them were used as herbal complements: *E. angustifolia, E. purpurea*, and *E. Pallida. Echinacea* contains chemical compounds liable for medicinal properties as: caffeic acid derivatives, polysaccharides, flavonoids, ketones, and lipophilic alkamides [27–34]. According to *in vitro* and *in vivo* studies, *Echinacea* showed effects on the cytokine production [35, 36], increasing the expression of CD69 [37], an impact on natural killer cells [38], as well as reducing illness severity, *in vivo*, as anti-inflammatory therapeutic agents on human monocytic THP-1 cells [39]. The latter is due to its efficacy on upper respiratory tract infections especially resulted from *E. purpurea* root derived a polysaccharide-containing water, and thereby activates phosphatidylinositol-3-kinase (PI3K) of following inhibition of TLR1/2 mediated stimulation of TNFα production [39]. Also, alkylamides and ketones of *Echinacea* extracts reported the anti-inflammatory effects [30, 40–42]. Another study in 2009 against H5N1 HPAIV strain showed that the extract of *E. purpurea* interferes with the viral entry into cells by blocking the receptor binding activity of the virus [43].

The application of medicinal plant extracts is still a controversial issue especially the plant extracts found in different manufacturing methods with varying use of plants parts and species. In this study, we evaluated the efficacy of *H. perforatum* and *Echinacea* as anti-viral agents against SARS-CoV-2 *in vitro*. The *H. perforatum* and *Echinacea* species applied in this study included aerial parts with high concentration of hypericin (main component 0.9 mg) and three different parts of type *E. purpurea* (root, seed, aerial parts) and one part of type *E. angustifolia* (root).

## Results

### Evaluation of Cytotoxicity of Hypericum perforatum and Echinacea

The cytotoxicity of *H. perforatum, Echinacea* and the *H.E.* mixture of both was evaluated on Vero E6 cell 48h post-treatment using MTT assay. The results showed that most of the plant extracts displayed minimal cytotoxicity at concentrations less than 12.5 µg/mL compared to the cell control (Figure 1). Also, the maximum non-toxic concentrations of *H. perforatum, Echinacea* and *H.E.* mixture were found to be: 1.56, 6.25, and 6.25 µg/mL; respectively.

**Figure 1.**
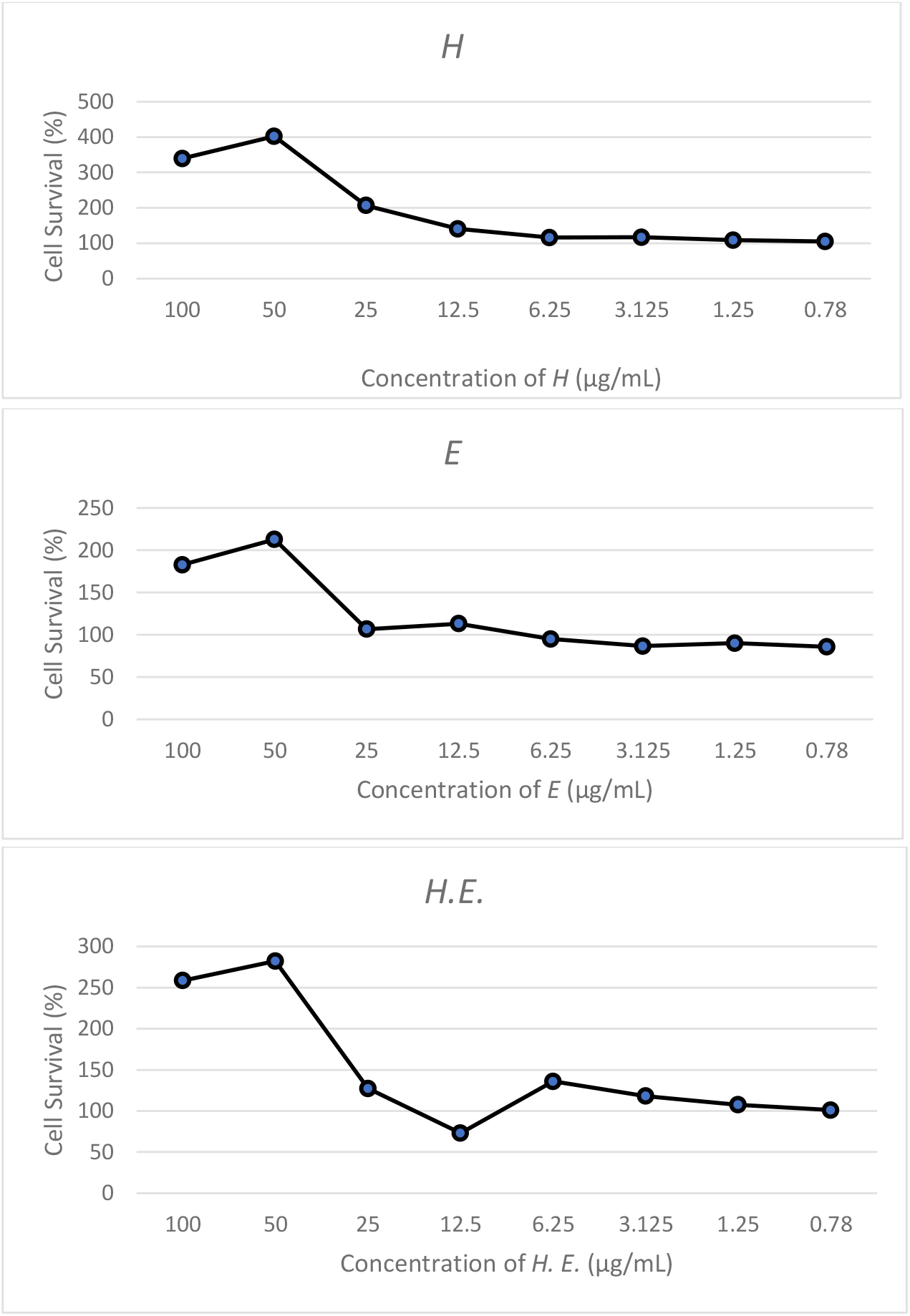
Cell viability of Vero E6 was determined by MTT assay. The survival rate of cells treated with different concentrations of *H. perforatum, Echinacea* and *H.E.* ranging from 100 to 0.78 µg/mL. The percentage cell viability was calculated relatively to cell control wells. More than 50% cell survival rate was considered to be the maximum non-toxic concentration of *H, E* and *H.E.* Statistical analysis showed that data were significant with *p*<0.005 (one-way ANOVA) of three independent experiments.

Serial dilutions ranging from 100-0.78 µg/mL were chosen for *in vitro* micro-inhibition assays, so, *H. perforatum* and *H.E.* showed IC50 values of 7.724 and 8.202 μg/mL, respectively, while the effect of *Echinacea* was not wasn’t concentration dependent as shown in Table 3 and Figure 2.

**Table 3.** Cellular toxicity and maximum non-toxic concentrations of *H. perforatum, Echinacea* and *H.E.* mixture against SARS-CoV-2 infection.

| Medicinal plant | $\text{IC}_{50}$ ($\mu\text{g/mL}$) | Max. non-toxic conc. ($\mu\text{g/mL}$) |
| --- | --- | --- |
| H | 7.724 | 1.56 |
| E | Nd | 6.25 |
| HE | 8.202 | 6.25 |

**Figure 2.**
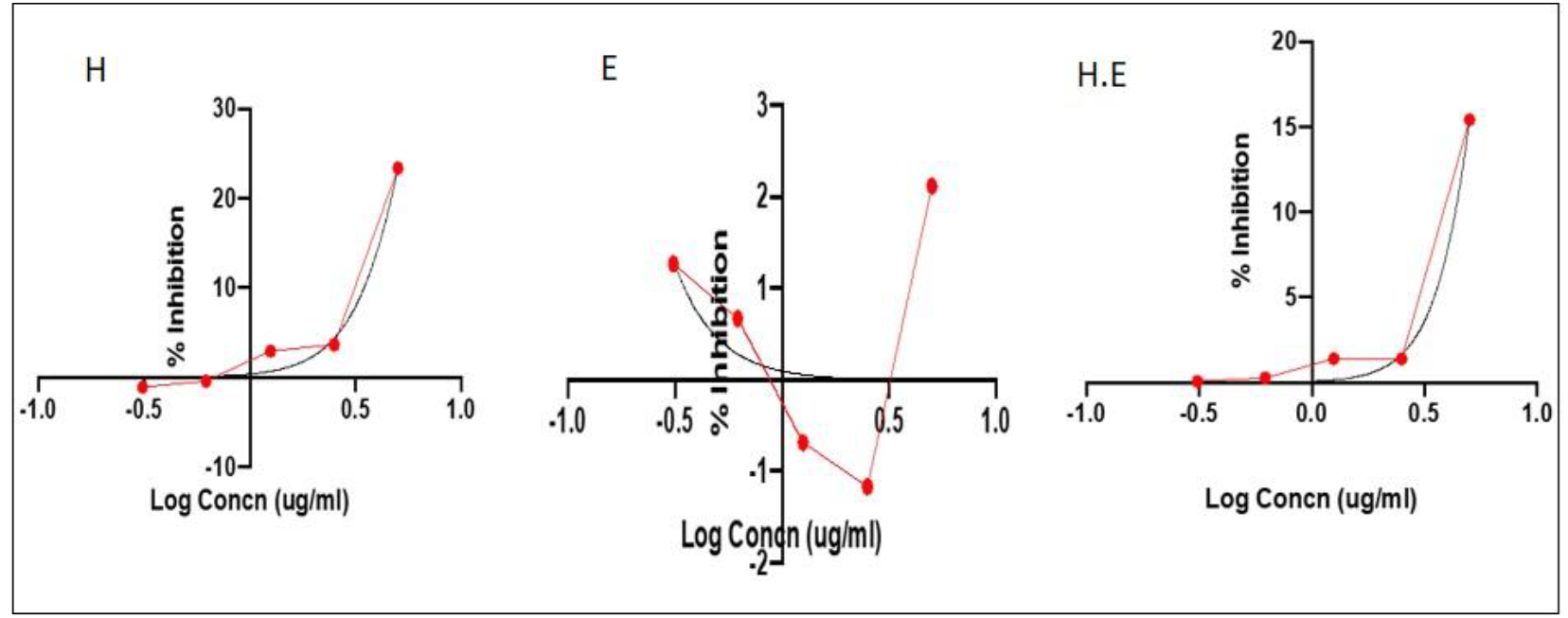
Dose-response curve analysis by Graph pad software for *H. perforatum, Echinacea* and *H.E.* activities. The black curve represents inhibition of SARS-CoV-2 infection (%) and the red dots represent log concentration (µg/mL).

### Anti-viral efficacy of Hypericum perforatum and Echinacea

The effect of the medicinal plants was evaluated by the anti-viral assays as following:

1. Direct treatment of virus–infected cells assay: In this assay the virus was incubated with cells for two hours at 37°C prior adding the medicinal plant mixture to the cells. Results from this assay showed that *H. perforatum* had the highest efficacy with IC50 of 1.56 μg/mL while *H.E.* mixture with IC50 of 6.25 μg/mL and the least effective was *Echinacea* of 6.25 μg/mL. Following the viral load with time (12, 18, 24, 36, 48hrs) after addition of the mixture showed a significant reduction of the viral load for *H. perforatum* up to 36hrs of addition (*p*=0.0047) followed by *H.E.* mixture up to 36hrs of addition (*p*=0.0048) and *Echinacea* up to 24hrs of addition (*p*=0.0060).
2. Pre-treatment of cells prior viral infection assay: In this assay the cells were incubated with medicinal plants for two hours at 37°C prior viral infection of the cells. The outcome of this assay showed that *H. perforatum* had an effective concentration of 1.56 µg/mL, while *H.E.* mixture had an effective concentration of 6.25 μg/mL and the least effective was *Echinacea* with 6.25 μg/mL concentration. Figure 4 showed the reduction in the viral load with *H. perforatum, H*.*E*. mixture and *Echinacea* over time (12hrs – 48hrs).
3. Virucidal activity assay: in this assay the virus was incubated with the medicinal plants for two hours prior addition to the cells. Results from this assay showed that *H. perforatum* had the highest effect of 1.56 µg/mL followed by the mixture *H.E.* of 6.25 μg/mL and the least effective was *Echinacea* of 6.25 μg/mL. The time of addition showed a significant reduction in the viral load for *H. perforatum* that lasted longer than 48hrs, and this result shown as well with *H.E.* mixture; and finally *Echinacea* displayed the best virucidal effect up to 36hrs.

The anti-viral effect of *H. perforatum, Echinacea* and *H.E.* (Figure 3 & 4) was obvious under maximum non-toxic concentration 1.56, 6.25, and 6.25 µg/mL, respectively, and the inhibition of *H. perforatum* on SARS-CoV-2 was the greatest from *H.E.* and *Echinacea* in the three anti-viral assays. Also, *Echinacea* was a weaker inhibitor than the *H. perforatum* and *H.E.* but slightly strong as a virucidal up to 24hrs.

**Figure 3.**
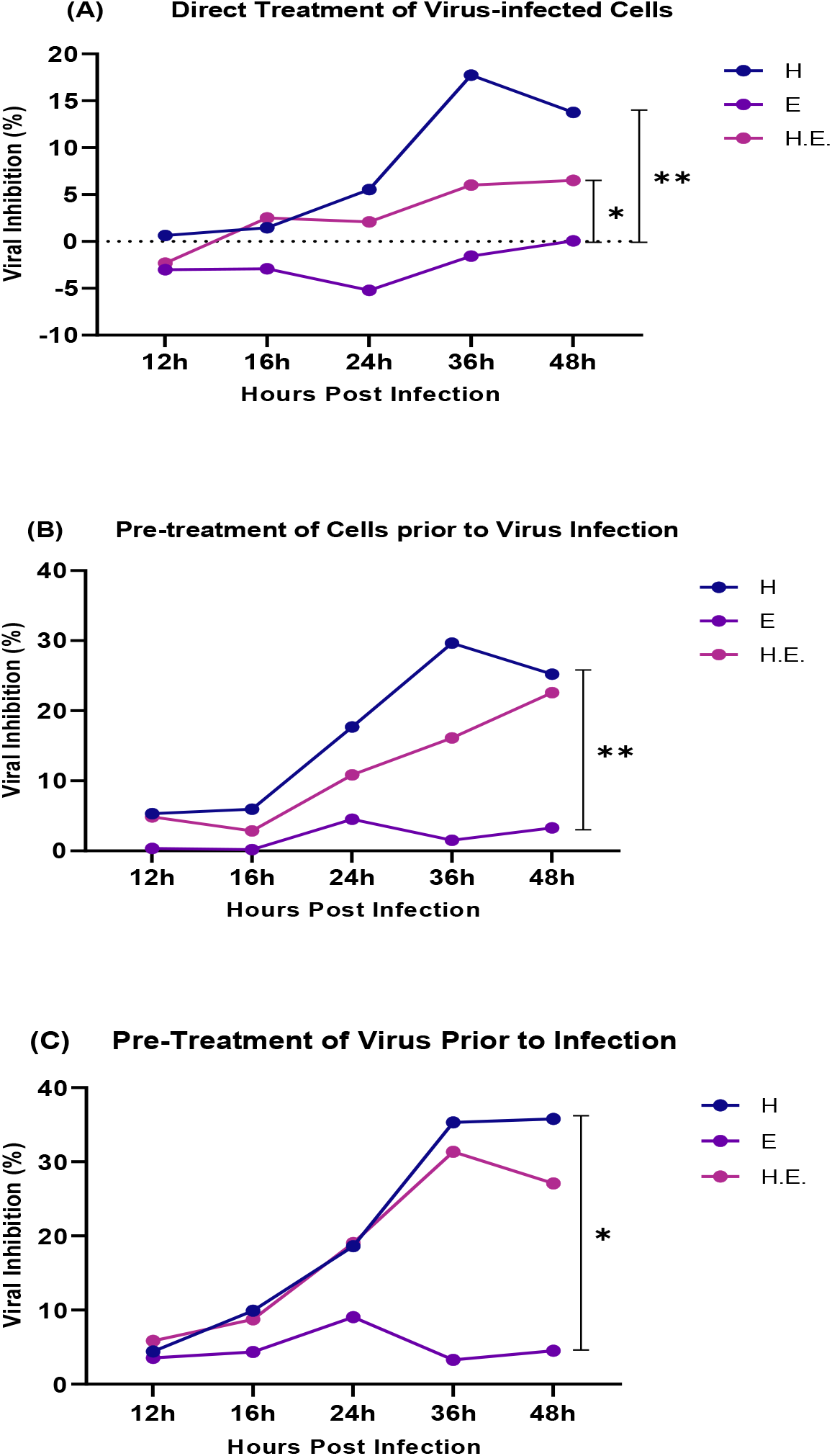
The inhibitory effect of *H. perforatum, Echinacea*, and *H.E.* on mRNA expression levels of SARS-CoV-2 RdRP gene in Vero E6 cells was evaluated by qRT-PCR. The impact of plant materials was detected by the three experimental designs: Direct Treatment of Virus-infected Cells (A), Pre-treatment of Cells Prior to Virus Infection (B), and Virucidal (C) assays; treated and infected cells were included as negative and positive controls, respectively. So, 100 TCID50 of the virus was added with the certain concentrations of the plant materials when the cells at 80% confluence were complete. The viral inhibition percentage was calculated relatively to virus control wells, and representatives of two independent experiments performed in triplicate are shown. Statistical analysis showed that data were significant with *p*< 0.05 and *p*< 0.005 (one- & two-ways ANOVA).

**Figure 4.**
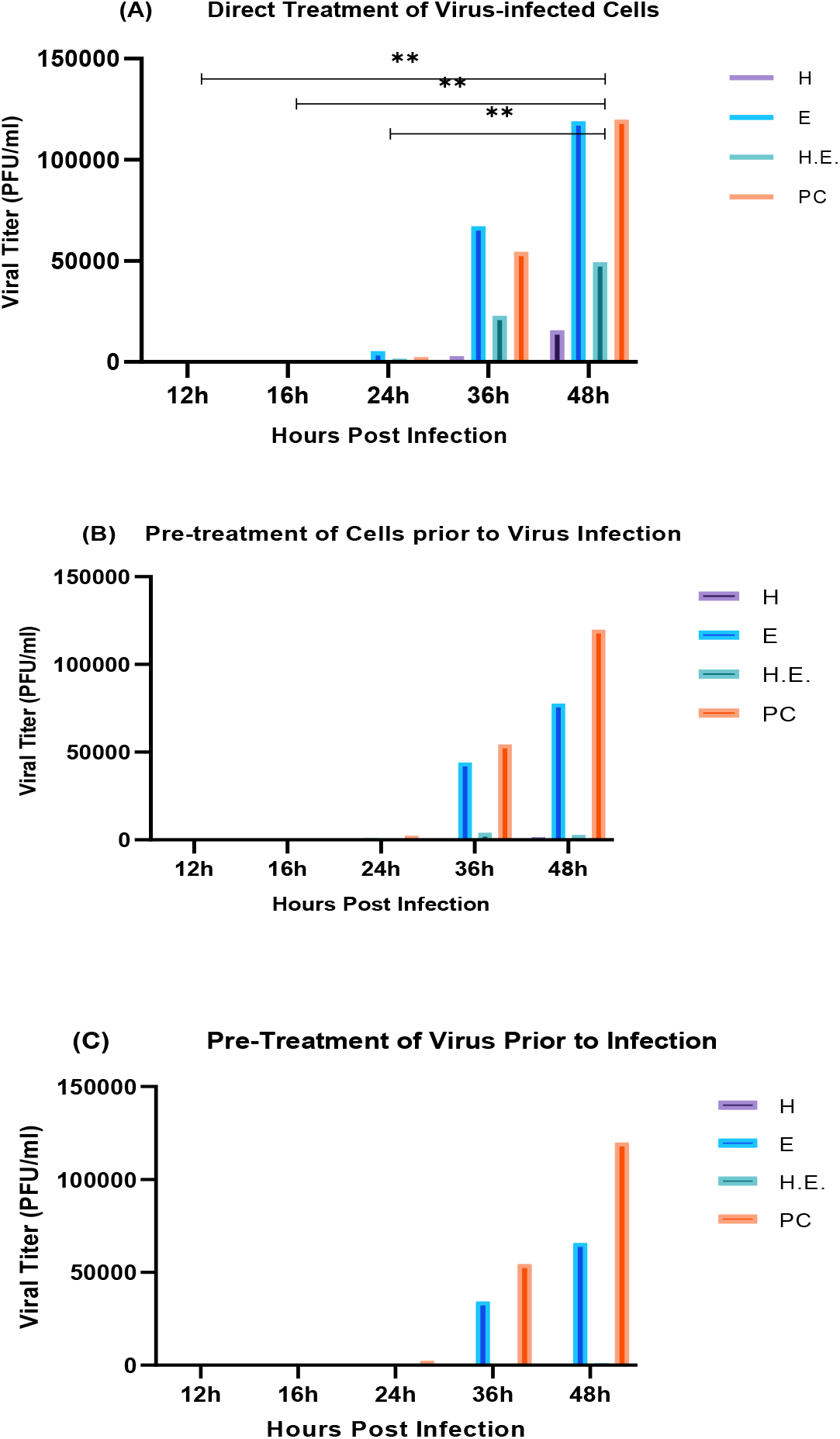
The effect of *H. perforatum, Echinacea*, and *H.E.* on viral load of SARS-CoV-2 infection in Vero E6 cells was evaluated by qRT-PCR. The impact of plant materials was detected by the three experimental designs: Direct Treatment of Virus-infected Cells (A), Pre-treatment of Cells Prior to Virus Infection (B), and Virucidal (C) assays; treated and infected cells were included as negative and positive controls (PC), respectively. So, 100 TCID50 of the virus was added with the certain concentrations of the plant materials when the cells at 80% confluence were complete. The effect of the plant materials was observed clearly with reduction in viral load at 1.56, 6.25, and 6.25 μg/mL, respectively. The viral load was calculated relatively to virus control wells, and representatives of two independent experiments performed in triplicate are shown. Statistical analysis showed that data were significant with *p*< 0.005 (one- & two-ways ANOVA).

## Discussion

COVID −19 pandemic is causing a global challenge to the world economic and healthcare systems. The pandemic is responsible for the death of over 1.8 million confirmed cases and 85.5 million infections worldwide (WHO). Although some vaccines are developed and are now released under emergency use because of the pandemic, the efficacy of the vaccines is still debatable specially with the emergence of new variants in the genomic structure [44–47]. Several non-specific treatment options were evaluated and entered clinical trial including the repurposing of known treatments against other diseases [48]. Therefore, improvement and investigation of an effective anti-viral therapy is a for treating SARS-CoV-2 infection.

In our study, the anti-SARS-CoV-2 effect of the medicinal plants *H. perforatum, Echinacea* and their combination was evaluated. The medicinal plants were purchased from a commercial source to ensure consistency of the composition and were tested either individually or combined together as a single treatment. Their mode of action was evaluated using three approaches namely direct treatment of virus–infected cells, pre-treatment of cells prior viral infection and virucidal approaches. The results were evaluated using qRT-PCR assay relative to the virus control (virus added to the cells without the medicinal plants) and were observed under the microscope for CPE effect on the cells and by colorimetric staining using crystal violet Figure 5.

**Figure 5.**
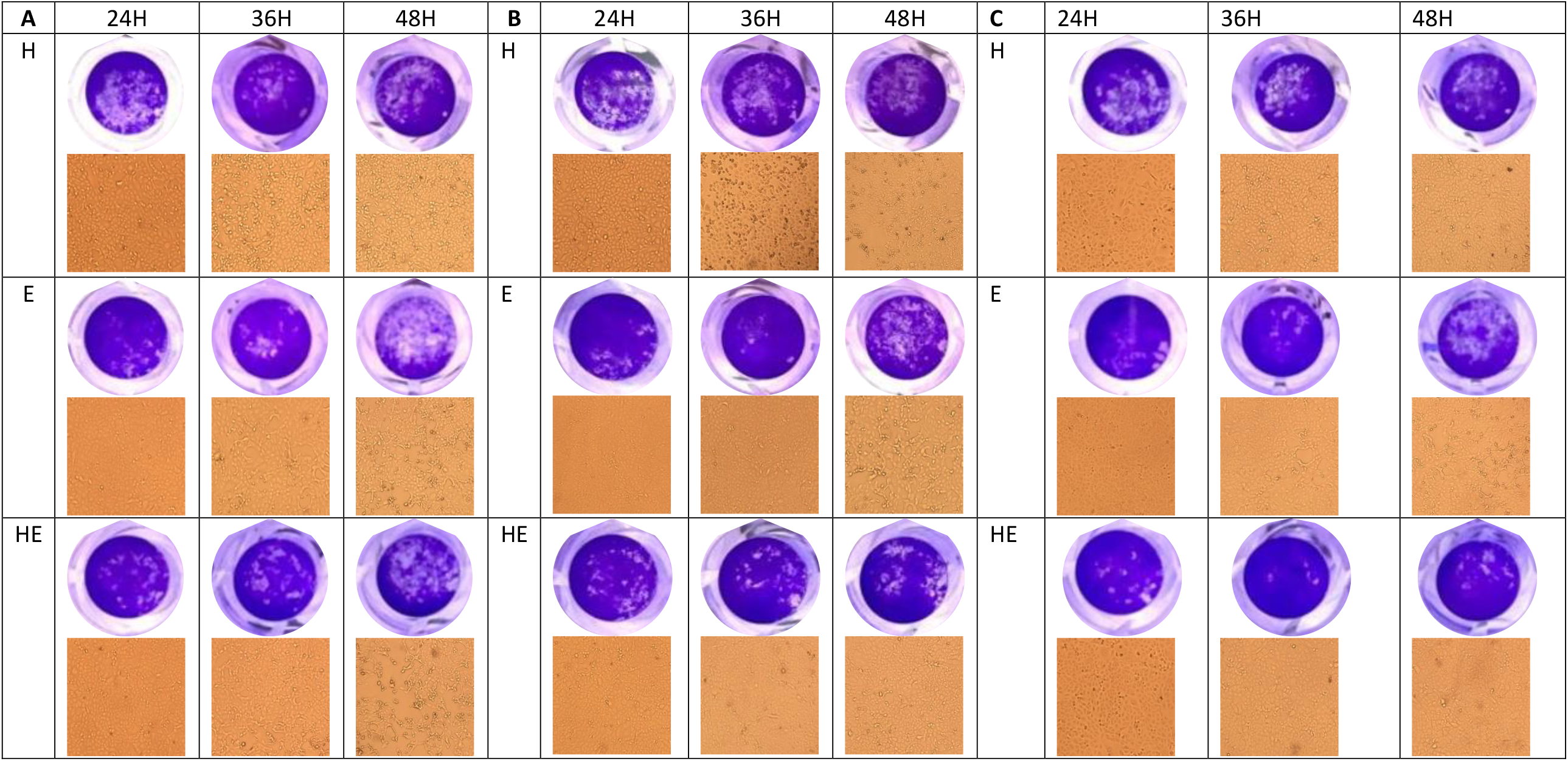
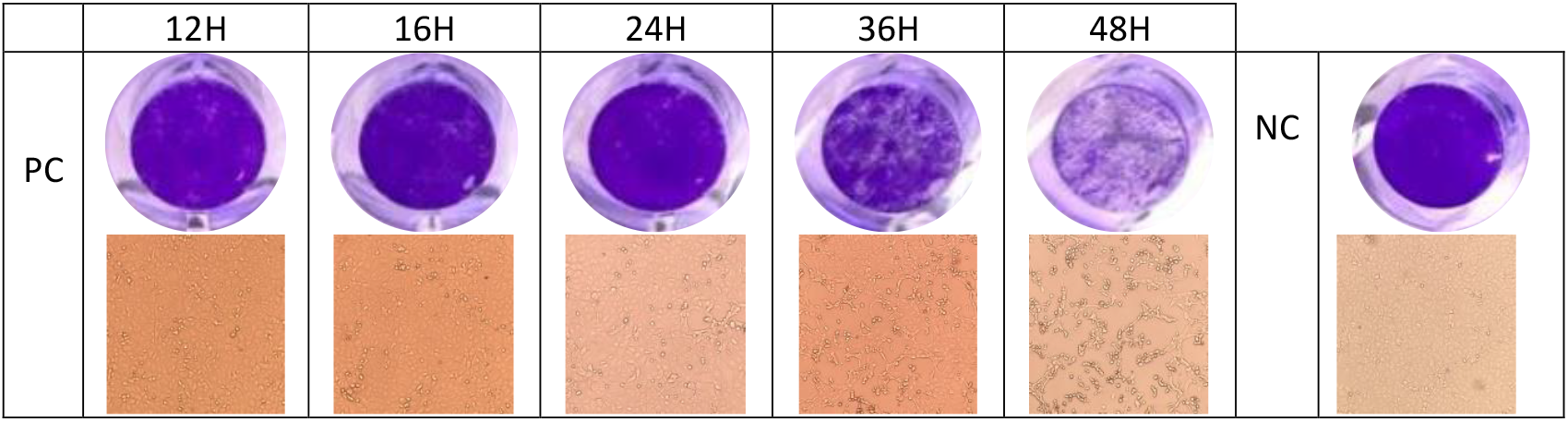
The impact of *H. perforatum, Echinacea* and *H.E.* on SARS-CoV-2 infected-Vero E6 cells by anti-viral activity essays. The cell viability and CPE through SARS-CoV-2 infectious cycle, from 24 to 48 hours post infection, were visualized by photographed images with a light microscope (magnification, x100) (orange squares) and by the crystal violet assay (blue circles). The images were captured when the CPE started on the positive control at 24hrs.

The cytotoxic effect of the tested medicinal plants was evaluated using MTT assay with results presented as percent of cytotoxicity relative to cell control (cells with no added tested medicinal plants). The IC50 of *H, E* and *H.E.* mixture was as follows 7.724, not detected and 8.202; respectively.

Our results demonstrated that the *H. perforatum*, containing 0.9 mg of hypericins, can significantly reduce SARS-CoV-2 viral load compared to the positive control (Table 4, Figure 3 & 4) through the viral infectious cycle from 0hr to 48hrs, and clearly this reduction in the anti-viral (treatment of virus-infected cells and pre-treatment of cells prior to virus infection) and virucidal assays.

**Table 4.**
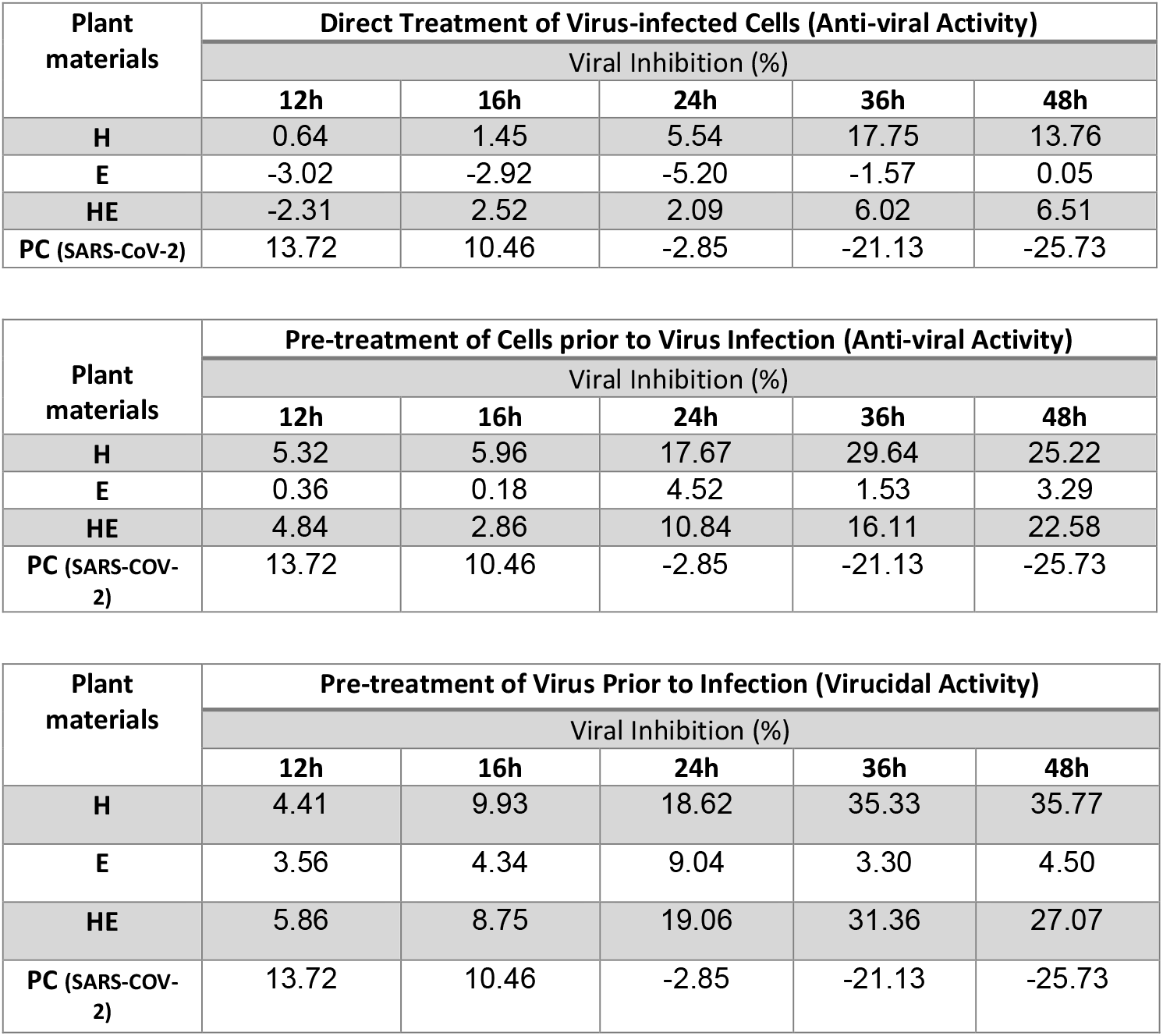
Inhibitory effects of *H. perforatum, Echinacea* and *H.E.* on SARS-CoV-2 mRNA expression level of RdRP gene through infection in Vero E6 cells.

Noticeably the results showed that the highest anti-viral effect was through the virucidal mechanism (Figures. 4 and 6) compared to the other two mechanisms with the highest inhibition observed at 36hrs after infection as shown in Figure 3. While, *H. perforatum* has the strongest inhibitory effect (35.77%) and reducing the viral load, up to 48hrs, compared to *Echinacea* and *H.E*. mixtur*e* that had 3.30 and 31.36%, up to 36hrs, respectively.

**Figure 6.**
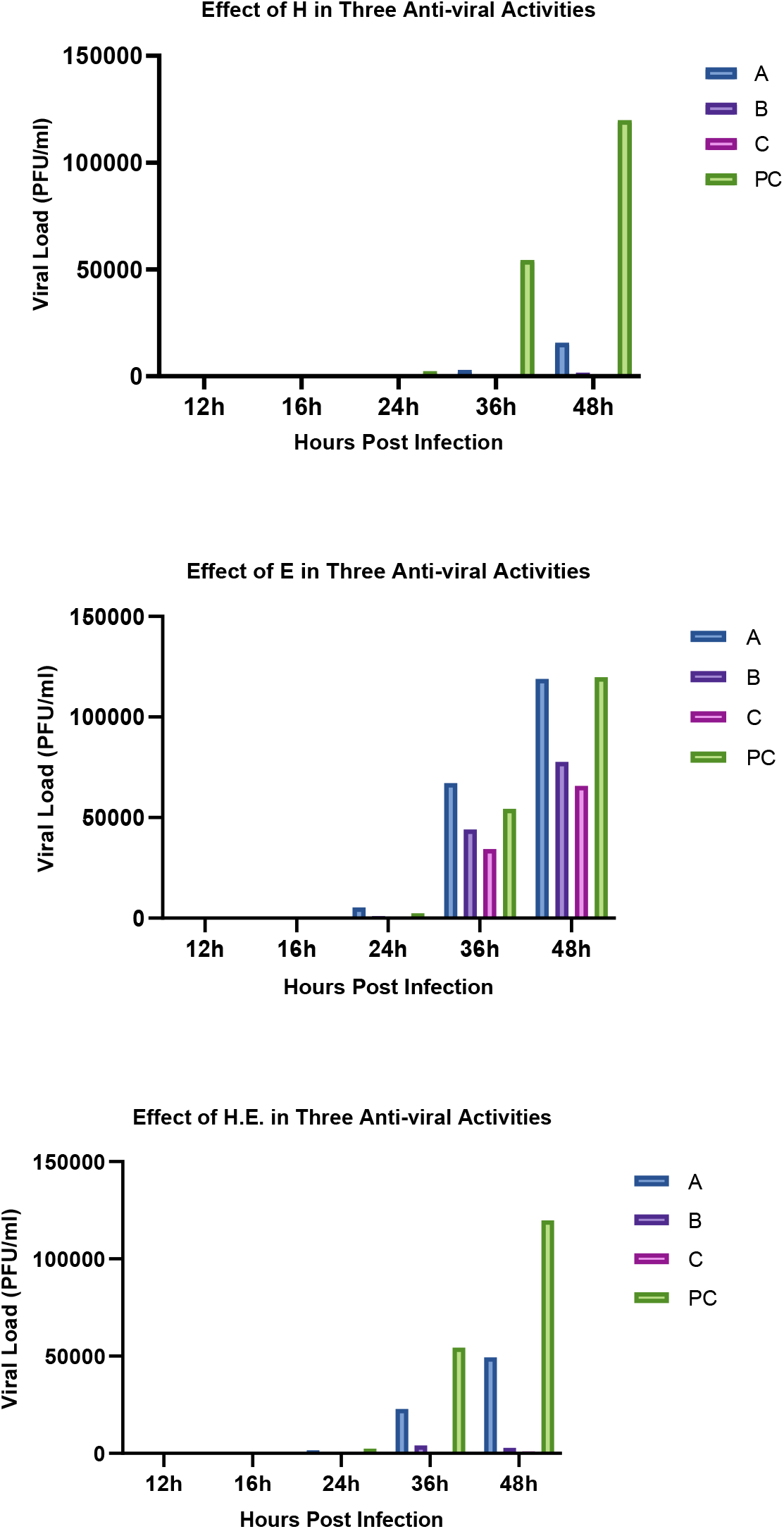
The effect of *H. perforatum, Echinacea*, and *H.E.* on viral load of SARS-CoV-2 infection in Vero E6 cells was evaluated by qRT-PCR. The impact of plant materials was detected by the three experimental designs: Direct Treatment of Virus-infected Cells (A), Pre-treatment of Cells Prior to Virus Infection (B), Virucidal (C) assays, and positive control (PC). The effect of the plant materials was observed clearly with reduction in viral load at 1.56 μg/mL for *H. perforatum*, then for *H*.*E* at 6.25 μg/mL, and finally for *Echinacea* at 6.25 μg/mL.

The anti-viral effect of the *H. perforatum* (with concentration of 1.56 μg/mL) was higher than *Echinacea* and *H.E.* mixture (with concentration of 6.25 µg/mL for both), and the effect was noted at lower IC50 and a higher reduction in viral load of SARS CoV-2. *H. perforatum* showed the highest inhibitory effect of in all three anti-viral assays Figure. 6, while *Echinacea* showed the lowest inhibitory effect. The *H.E.* mixture showed lower inhibition than *H. perforatum* alone that would be expected as the active compounds were diluted by the addition of *Echinacea*.

Hypericins were reported to be the active anti-viral compounds in *H. perforatum* extract against IBV *in vitro* and *in vivo* [14]. The *H. perforatum* ethylacetate (HPE) extract includes: hyperoside, quercitrin, quercetin, pseudohypericin, and hypericin. The anti-viral effect of HPE was reported previously against bovine diarrhea virus (BVDV) *in vitro* [48] and hepatitis C virus (HCV) *in vivo* [49] with hypericin of *H. perforatum* extract defined as the active ingredient. Other studies in 2001, 2009, and 2012 showed that *H. perforatum* extract had an anti-viral effect against influenza A virus and HIV [50–52], suggesting that HPE could be developed and used as an anti-viral drugs. In addition, the anti-viral effect of *H. perforatum* extract and HPE had a remarkable decrease in the concentration of IL-6 and TNF-α through the NF-κB in lung tissue of mice infected with an influenza A virus [52] and in the trachea and kidney for chickens infected with IBV, mainly from hypericin content [14].

*Echinacea*, as indicated by the supplier insert is composed of two types (*E. purpurea* (root, seed, aerial parts) and *E. angustifolia* (root)). The beneficial effects of *Echinacea* were reported previously in both *in vitro* and *in vivo* studies as preventive and treatment for respiratory tract viruses due to its active components such as: polysaccharides, glycoproteins, caffeic acids, and alkamides [53–56]. Also, *Echinacea* was reported to achieve the inhibition against influenza A and B, respiratory syncytial virus (RSV), parainfluenza virus, and herpes simplex virus [41, 57]. *E. purpurea* was also demonstrated to interfere with H5N1 and HPAIV cellular entry into cells by blocking the receptor binding activity of the virus, and to emphasize the direct contact with it and the virus in order to gain the maximum inhibition of replication [41]. Because of *Echinacea’s* reduction on virus cytokine liberation, it can shorten and relieve the severity of symptoms as a result of pro-inflammatory cytokines release [58, 59]. An *in vivo* study is needed to evaluate the effect of *Echinacea* on cytokine regulating during COVID-19 infection together with the anti-viral effect found in this study of *H. perforatum*. Lately, Signer and his group showed *in vitro* that Echinaforce^®^, type of *E. purpurea* and from a commercial product, is virucidal against human coronavirus (HCoV) 229E, MERS- and SARS-CoVs, as well as SARS-CoV-2 [60]. In hence, gaia HERBS^®^ *Echinacea* has consistent result as an effectively prophylactic treatment in virucidal by blocking the receptor binding activity of the virus; and anti-viral activities (pre-treatment of cells prior to virus infection) by inhibition of the viral load, *in vitro* against SARS-CoV-2 infection.

An individual finding of our study, the application of both *H. perforatum* and *Echinacea* (100 mg/mL, of each) exerted the high inhibition in the two anti-viral and virucidal activities against SARS CoV-2 infection cycle up to 48hrs (Table 4, Figure 3). So, this result provides, for the first time, clear evidence that the mixing materials of *H. perforatum*, containing pseudohypericin, hypericin, hyperforin, adhyperforin, quercetin, quercitrin, and other components, possess anti-SARS CoV-2 activities with the components of *Echinacea* (Table 2). In addition, the mixture effects correlated with reducing the viral load up to 48hrs post infection (Figure 6).

**Table 2.** Components found in *Hypericum perforatum* aerial part and *Echinacea* species. 412

| *Hypericum perforatum* | *Echinacea* species |
| --- | --- |
| Aerial parts:<br>naphthodianthrones<br>(hypericin and<br>pseudohypericin),<br>acylphloroglucinols<br>(hyperforin and adhyperforin),<br>flavonol glycosides<br>(quercetin, quercitrin,<br>isoquercitrin, rutin,<br>hyperoside, kampferol,<br>leteolin, myricetin),<br>biflavones<br>(<sup>13,118</sup>-biapigenin, <sup>139,118</sup>-<br>biapigenin (amentoflavone)),<br>Proanthocyanidins<br>(procyanidin B2) and<br>phenyl propanes<br>(chlorogenic acid and caffeic<br>acid) [65, 66] | *Echinacea angustifolia* root [63]<br>Essential oil<br>Polyacetylenes<br>Alkamides<br>Alkaloids<br>Caffeic acid derivatives<br>Polysaccharides and glycoproteins<br>*Echinacea purpurea* root [63]<br>Essential oil<br>Alkamides<br>Alkaloids<br>Caffeic acid derivatives<br>Polysaccharides and glycoproteins<br>*Echinacea purpurea* seed [64]<br>Essential oil<br>Fatty acids [64]<br>*Echinacea purpurea* aerial parts [63]<br>Essential oil<br>Alkamides<br>Flavonoids and anthocyanins<br>Caffeic acid derivatives and phenolic acids<br>Polysaccharides |

For clinical trial, the mixture may empower the inhibition of the virus by up-regulating the mRNA expression process, lower the viral load, and neutralizing the virus envelop receptor as anti-viral or/and virucidal activities, respectively; and definitely it is related to the pro-inflammatory cytokines such as: IL-6, TNF-α, INF-β as anti-inflammatory therapy. It can be useful to precede and develop a new technique for treatment SARS-CoV-2 patients in mild or sever cases, protecting people who may contact infected patients, or/and having positive COVID-19 test. In addition, regarding the using of mixture materials in clinical trial, it was clear that preparation 100 mg/mL of each medicinal plant was safe *in vitro* rather than 200 mg/mL in testing the efficacy against SARS CoV-2 infection, which was recommended using *H. perforatum* and *Echinacea* for patients in different-time administration. Alternatively, *H. perforatum* is preferable single used as anti-viral treatment for the mild to sever case-patients because it demonstrated a high efficacy against SARS-CoV-2 infection. On the other hand, either *Echinacea* or *H.E.* is suggested to be admitted as prophylactic treatment with considering the dose, side effects, drug interactions, and toxicity.

## Materials and Methods

### Cell Cultures

Vero E6 cells (ATCC^®^ CRL-1586™) were maintained in 75 cm2 cell culture flask containing Dulbecco’s modified Eagle’s medium (DMEM) supplemented with 10% fetal bovine serum (SIGMA) at 37 °C in an incubator containing 5% CO2. For preparation 96-well plates with Vero E6 cells, the plates were seeded with 3.10^4^ cells/well and incubated for 24hrs at 37°C with 5% CO2 until a confluent monolayer was attained [61].

### Viral isolation and stock

The procedure of viral isolation of our human SARS-Cov 2 clinical patient isolate (SARS-CoV-2 /human/SAU/85791C/2020, gene bank accession number: MT630432) was described in [62]. The viral production was done in 75 cm^2^ cell culture flask containing Vero E6 cells in Minimum Essential Media (Gibco, ThermoFischer) (MEM) with 4% of fetal bovine serum and 1% glutamine. Cytopathic effect was monitored daily under an inverted microscope. In approximately 72hrs, nearly complete cell lysis was observed of the TCID50 of the strain at 3.16.10^6^ infectious particles per mL. The viral supernatant was used for inoculation on 96-well plate.

### Commercial Products of Hypericum perforatum and Echinacea Extracts

*H. perforatum* and *Echinacea* products were purchased from gaia HERBS^®^ as gelatin capsules and as liquid; respectively. Table 1 shows the full the information provided on the packaging of products along with information leaflets. The products were sold as dietary and herbal supplements carried the Certified B Corporation logo which transmits a level of quality (Table 1, 2).

**Table 1.** The products materials used from gaia HERBES^®^.

| Proprietary Extract Blend | Amount per serving | ĖHerb Equivalency |
| --- | --- | --- |
| St John's Wort (*Hypericum perforatum*) aerial parts, St John's Wort (*Hypericum perforatum*) aerial parts extract. | 500 mg | Hypericins (0.9 mg) |
| Organic *Echinacea purpurea* root, Organic *Echinacea angustifolia* root, Organic *Echinacea purpurea* seed, Organic *Echinacea purpurea* Aerial parts. | 2 mL | 444 mg/mL |

### Sample Preparation

Gelatin capsules were separated and the contents emptied into a weighing boat to ensure the coating was not included. Then, the plant materials, capsule sample and liquid, in concentration of 100 mg/mL [67] were dissolved in Dimethyl sulfoxide (DMSO, ≥99.5%, plant cell culture tested, SIGMA) and filtered into 0.22 µm [68]. All the extracts were filtered to remove any plant fibers. The mixture (*H.E.*) of *H. perforatum* and *Echinacea* was prepared by mixing each plant material at 100 mg/mL. Finally, the stocks were stored at −20 °C.

### Cytotoxicity Assay

The cytotoxicity assays were performed by 3-(4,5-dimethylthiazol-2-yl)-2,5-diphenyltetrazolium bromide (MTT, Roche, ref:11465007001) protocols according to the previous study [47] with few modifications. Vero E6 cell monolayers with 4.10^3^ cells/mL plated onto 96-well culture plates were washed 3 times with phosphate buffer saline (PBS) 1x, pH 7.4. 100 µl of prepared *H. perforatum, Echinacea* and *H.E.* diluted in serum-free DMEM medium (two-fold dilutions, ranging from 0.78-100 µg/mL) and another 100 µl of growth medium added onto the wells (3 wells per each dilution). Control cells were incubated with the same corresponding concentrations of DMSO solutions as negative controls. The cells were cultured at 37°C for 48h in a CO2 atmosphere, and the supernatant was taken out and washed 3 times with PBS, then 20 μL of MTT solution was added to each well and incubated for 4hrs at 37°C. Solubilization solution was added and incubated for overnight at 37°C. Then, the evaluation of the plate was done using a reader (Synergy 2 microplate, BIOTIK) with a reference wavelength of 570 nm, and the OD570 was recorded. Cell survival rates were calculated from mean values according to the following equation: (OD570 drug)/ (OD570 control) × 100%.

### In vitro Micro-inhibition Assay

According to a method described previously [69] and with some modifications, the plant materials activity against SARS-CoV-2 was evaluated from the curve that was generated by plotting percentages of virus inhibition against concentrations of plant materials. Briefly, 96-well plates were prepared as above mentioned with Vero E6 cells. Then, the cells were washed twice with PBS, and two-fold serial dilutions of plant materials (*H. perforatum, Echinacea* and *H.E.*) (0.78-100 µg/mL) in medium were challenged with 100 TCID50 of SARS-CoV-2 and incubated for three days at 37°C/5% CO2, the results were quantified as previously described. IC50 was expressed relative to the virus control using dose-response curves; and, in Graph Pad software for anti-viral activity curves analysis, a trendline that best suited the curve was selected and the correlating equation was used to calculate IC50 values [69].

### Anti-viral Activity Assays

The effect of the medicinal plants was tested by the anti-viral assays as following:

### 1. Direct Treatment of Virus-infected Cells (Anti-viral Activity)

To determine the effect of *H. perforatum, Echinacea* and *H.E.* on SARS CoV-2 infected cells, the Vero E6 cells cultured on 96-well plates were infected with 100 TCID50 of the virus for 2hrs at 37°C, as described previously [14]. Then, treated with 1.56, 6.25, and 6.25 µg/mL of *H. perforatum, Echinacea* and *H.E.*, respectively, at 37°C for 48h. The supernatant of the cell samples was collected at every 12, 16, 24, 36, and 48hrs (3 wells per each time) [70] for qRT-PCR [62]. According to above description, the relative mRNA expression levels of SARS-CoV-2 RdRP gene was detected, the viral load of the cell samples was determined, and the level of virus neutralization was assessed too. SARS-CoV-2-infected Vero E6 cells were processed as controls.

### 2. Pre-treatment of Cells Prior to Virus Infection (Anti-viral Activity)

To investigate direct effects of *H. perforatum* on the virus, the Vero E6 cells cultured on 96-well plates were treated with 1.56, 6.25, and 6.25 µg/mL of *H. perforatum, Echinacea* and *H.E.* solutions, respectively, at 37°C for 2hrs, and then washed 3 times with PBS solution [14]. The cells were subsequently infected with SARS-CoV-2 (100 TCID50) and cultured at 37°C for 48hrs. Also, the supernatant was collected every 12, 16, 24, 36, and 48hrs (3 wells per each time) [70] for qPCR. We extend the observation till 48hrs to assess the extracts efficacy after 36hrs post-infection. The relative mRNA expression levels of SARS CoV-2 RdRP gene, the viral load, and the level of virus neutralization were assessed with qRT-PCR [62].

### 3. Pre-Treatment of Virus Prior to Infection (Virucidal Activity)

To analyze the impact of *Echinacea* on cell whether to inhibit or not SARS-CoV-2 infection through neutralization binding receptor of the virus, 100 TCID50 SARS-CoV-2 was incubated with 1.56, 6.25, and 6.25 µg/mL *H. perforatum, Echinacea* and *H.E.*, respectively, at 37°C for 2h [14]. The three extract-treated SARS-CoV-2 were subsequently used to infect Vero E6 cells at 37°C for 48hrs. As described above, the relative mRNA expression levels of SARS CoV-2 RdRP gene, viral load, and the level of virus neutralization were assessed with qRT-PCR [62] of every 12, 16, 24, 36, and 48hrs supernatants cell samples [70]. During the post infection cycle, the plates were analyzed by observing virus-induced CPE by light microscope (Nikon-ECLIPSE-T*i*), and plaque formation was determined by crystal violet staining (C0775; Sigma-Aldrich), as previously described [47]. The supernatants were collected and further used for qRT-PCR.

### Quantitative Real-Time PCR (qRT-PCR)

Viral RNA was extracted from all samples collected directly from the 96 well plates of the anti-viral assay, as previously described [62] using the QIAmp Viral RNA Mini Kit (Qiagen, Germany) according to the manufacturer’s instructions. Relative quantification of the SARS-CoV-2 viral load was performed by one-step dual-target real time RT-PCR (RealStar SARS-CoV-2 RT-PCR Kit 1.0, Altona Diagnostics, Germany) according to the manufacturer’s instructions using a 7500 Fast Real-Time PCR System (Applied Biosystems, U.S.A.). The PCR detects a beta-coronavirus specific target (E-gene), a SARS-CoV-2 specific target (RdRP-gene) and an internal control. The decrease in viral load was expressed by comparing the cycle threshold (CT) values from each sample relative to the CT values of the pretreatment inoculated sample (with the SARS-CoV-2 specific primers). The SARS-CoV-2 titers were expressed as PFU equivalents per mL (PEq/mL) using a standard curve (standard: serial dilutions of the viral stock) and choosing dilutions of the original sample (10-1 to 10-8) with CT values in the exponential phase. Each run included a positive viral template control and no-template negative control. Each sample was tested in duplicate and the mean is reported as PEq/mL.

## Statistical analysis

Data was analyzed with one- or two-way ANOVA with a Tukey’s test for multiple comparisons. *P* < 0.05 and < 0.005 are considered statistically significant. All analyses were performed with GraphPad Prism, version 8.

## Acknowledgments

The authors would like to acknowledge the technical support of Mr. Ahmed M Hassan and Dr. Ahmed M Tolah for preparing the virus isolate and virus stock in the Biosafety level-3 lab of the Special Infectious Agents Unit, King Fahd Medical Research Center, King Abdulaziz University.

